# Cerebellar control over inter-regional excitatory/inhibitory dynamics discriminates execution from observation of an action

**DOI:** 10.1101/2024.05.21.595114

**Authors:** Roberta Maria Lorenzi, Gökçe Korkmaz, Adnan A.S. Alahmadi, Anita Monteverdi, Letizia Casiraghi, Egidio D’Angelo, Fulvia Palesi, Claudia A.M. Gandini Wheeler-Kingshott

## Abstract

The motor learning theory anticipates that cerebro-cerebellar loops perform sensorimotor prediction thereby regulating motor control. This operation has been identified during action execution (AE) and observation (AO) but the causal interaction between the cerebellum and cerebral cortex remained unclear. Here we used Dynamic Causal Modelling (DCM) to study functional MRI (fMRI) data obtained during a squeeze ball task in either the AE or AO conditions. In both cases, active regions included bilateral primary visual cortex (V1), left primary motor cortex (M1), left supplementary motor and premotor cortex (SMAPMC), left cingulate cortex (CC), left superior parietal lobule (SPL), and right cerebellum (CRBL). AE and AO networks showed the same fixed effective connectivity, with pathways between V1, CRBL, SMAPMC and CC wired in a closed loop. However, the cerebellar communication towards the cerebral cortex switched from excitatory in AE to inhibitory in AO. Moreover, in AE only, signal modulation was non-linear from SMAPMC to CRBL and within the CRBL self-connection, supporting the role of the CRBL in elaborating motor plans received from SMAPMC. Thus, the need for motor planning and the presence of a sensorimotor feedback in AE discriminate the modality of forward control operated by the CRBL on SMAPMC. While the underlying circuit mechanisms remain to be determined, these results reveal that the CRBL differentially controls the excitatory/inhibitory dynamics of inter-regional effective connectivity depending on its functional engagement, opening new prospective for the design of artificial sensorimotor controllers and for the investigation of neurological diseases.

## 1 Introduction

Motor tasks trigger an intense neuronal activity between brain areas engaging multiple cortico-cerebellar loops. Theory anticipates that the cerebellum (CRBL) plays a key role in motor control and planning as well as in cognitive processing, whereby it performs error detection and computes functional predictions^1^. To perform a task, the CRBL is thought to use cortical instructions for calculating the inverse model needed for the action, which is then transferred back to cortical regions for its execution. This sensorimotor control process operates in forward mode and makes the study of cerebro-cerebellar loops fundamental for understanding brain function and dynamics^2,3^.

Functional magnetic resonance imaging (fMRI) protocols have recently been developed to investigate how motor tasks shape the recorded Blood Oxygen Level Dependent (BOLD) signal in different brain regions. The effect of grip force (GF) was studied in squeeze ball visuomotor tasks during action execution (AE) and action observation (AO, i.e., subjects observed a video of the same task rather than actually squeezing the ball)^4,5^. Results of task performance shed light on how sensorimotor processes are physiologically integrated and are affected by the presence of neurological diseases (for example, the same AE task showed an altered BOLD-GF relation in multiple sclerosis, MS)^6^. During AE and AO, cerebellar and cortical areas involved in motor planning, motor execution, visual processing, and associative functions, were activated with different modulations of the BOLD signal by the applied GF. Notably, the primary motor cortex showed a linear positive BOLD-GF relation, while cortical areas involved in motor planning and motor control showed both linear and non-linear BOLD-GF relations^5^. Despite these studies, what remains unknown is the causal influence between pairs of active regions, which is essential to understand network interactions in AE and AO and to gain insights on role that the CRBL plays in forward control loops. Moreover, the analysis of the shape of the BOLD-GF relation measured at the macroscale level must reflect microscale neuronal phenomena, so that non-linearities in the system inform about the cellular mechanisms involved. Indeed, for example, single cell recordings demonstrated that capillary vasodilation in response to different frequencies of neuronal stimulation is non-linear in the cerebellar vermis and hemispheres^7^. Bridging the gap between macro and micro scale is therefore essential^8^.

Recently, neuroinformatic frameworks have offered the opportunity to investigate brain dynamics on multiple scales by integrating computational models of neuronal activity with ensemble recordings of brain function (such as BOLD signals)^8–10^ In particular, modelling revealed that the cerebellum could control cortical dynamics and perform forward control operations^2,3^. In this context, Dynamic Causal Modelling (DCM) represents a convenient framework that allows us to link neuronal activity to the BOLD signal, estimating the causal influence between regions involved in tasks such as AE and AO^11^. DCM convolves a neural model with an haemodynamic response function to estimate the BOLD signal response generated by neuronal mechanisms; its mathematical framework relies on Bayesian inference^12^, which consists in updating parameter priors at the neuronal level, by iteratively comparing a simulated BOLD signal with the observed data^11,13^. The output, called *effective connectivity*, quantifies the effect of one brain region onto another for a given task. Effective connectivity is computed as the sum of a fixed effect (reporting whether two regions are causally connected or not) and a modulation term (reporting whether connectivity is modulated by the task and how). Moreover, DCM indicates the connection directionality and measures its excitatory/inhibitory nature and strength.

In this study we have used DCM to investigate the causal interaction between regions involved in AE and AO during a visually guided squeeze ball task. We assessed effective connectivity between critical brain areas, including the bilateral primary visual cortex (V1), the left primary motor cortex (M1), the left supplementary motor and premotor cortex (SMAPMC), the left cingulate cortex (CC), the left superior parietal lobule (SPL), and the right cerebellum (CRBL). From this study the CRBL emerges as a key region that not only intervenes in the execution of a motor plan but that it is essential for predicting and updating the plan in forward mode. The CRBL excites SMAPMC through a non-linear modulation in AE but, strikingly, it inhibits SMAPMC through a linear modulation in AO, probably because the sensory feedback is not present and there is no need for an inverse model of the motor action. This study supports the forward control theory of the CRBL implying different mechanisms depending on the task in which the cortico-cerebellar loops are engaged.

## 2 Results

The fMRI recordings of subjects engaged in AE and AO squeeze ball tasks were analysed using DCM to quantify the causal dependencies between nodes through the identification of BOLD modulation and directional differences. Multiple network models were hypothesised and then scored with Random Effect Analysis Bayesian Model Selection (RFX-BMS). We first considered network models with fixed effective connectivity (Step1) and then we introduced modulation on the fixed effective connectivity (Step2). (See Methods Figure 4 and 5 for details).

### 2.1 Fixed Effective Connectivity (Step1)

RFX-BMS identified a specific network model as the winning model with a probability > 90% of explaining the observed data better than other models for both AE and AO (Figure 1A.1, A.2). This model indicates that the driving region V1 is connected to motor areas both for planning (CRBL and SMAPMC) and execution (M1). From the Step1 winning model, it emerged that there is a clear visuo-to-plan functional loop, including bidirectional connections between V1, CRBL, SMAPMC, and CC plus a cerebellar self-connection; M1 sits right in the middle of this visuo-to-plan loop, with bidirectional connections to all three regions (CRBL, SMAPMC and V1) plus bidirectional connections to SPL (Figure 1A.3).

**Figure 1.**
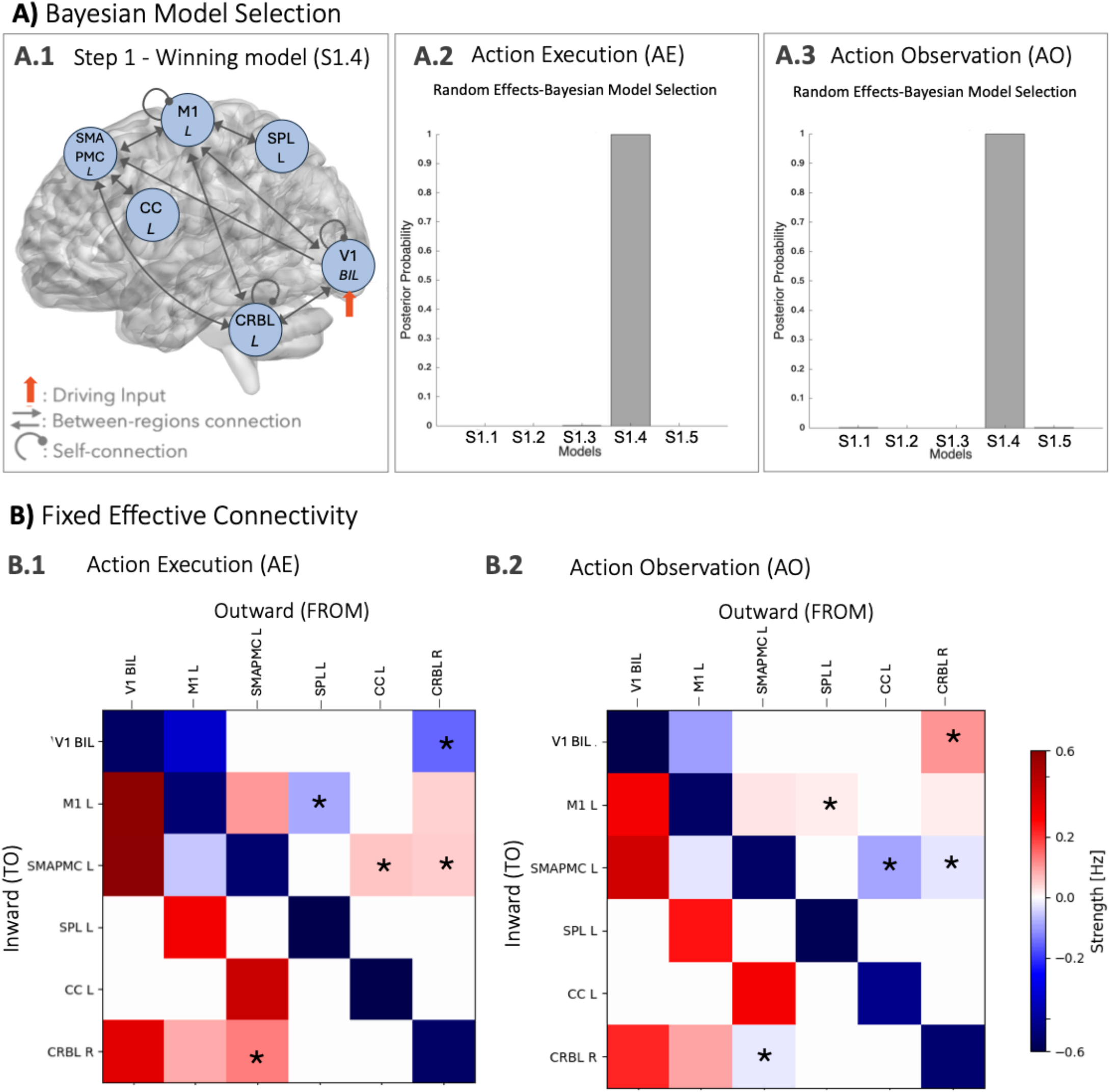
Fixed Effective Connectivity of the visuomotor network. **A)** Bayesian Model Selection with random effects (RFX-BMS) identifies Model S1.4 (Panel A.1) as the “winning model” with posterior probability > 90% both in action execution (AE in panel A.2) and action observation (AO in panel A3). **B)** Effective connectivity strength in Hz (see eq. 1 - Methods) with positive (red) values for excitatory connections and negative (blue) values for inhibitory connections. White values correspond to extrinsic connections not included in the model. Self-connectivity (diagonal) cannot be null by definition (see Methods) and it shows a remarkable intrinsic activity of all the regions involved in both AE and AO (AE in panel B1, AO in panel B2). The strongest excitatory connections are from V1 to SMAPMC, both in AE and AO, and from V1 to M1 only in AE. Connections that present a different excitatory/inhibitory nature in AE and AO are highlighted with a *; note that the crossed connections CRBL-R→SMAPMC-L and SMAPMC-L→CRBL-R are excitatory in AE and inhibitory in AO. V1 = bilateral primary visual cortex, M1 = left primary motor cortex, SMAPMC = left supplementary motor and premotor cortex, CC = left cingulate cortex, SPL = left superior parietal lobule, CRBL = right cerebellum.

The fixed effective connectivity strength (Hz) and its excitatory/inhibitory nature for each pair of regions are represented in Figure 1 B.1 and B.2 for AE and AO, resulting in non-symmetric connection matrices between cortical regions and the CRBL. Self-connectivity of all regions (reported on the diagonal) turned out to be inhibitory with a comparable strength for both AE and AO. Extrinsic (i.e., between-region) connectivity showed a predominance of excitation in both AE (73%) and AO (66 %) (Figure 1B.1, B.2). The strongest excitatory connections were from V1 to SMAPMC both in AE and AO, and from V1 to M1 in AE. A number of changes were observed between the excitatory and inhibitory nature of connections between some regions in AE or AO (highlighted with * in Figure 1.B): 1) SPL to M1 was inhibitory in AE and excitatory in AO; 2) SMAPMC inward and outward connections were different in AE compared to AO: the fixed effective connectivity from CC to SMAPMC was excitatory in AE but inhibitory in AO; 3) The most striking changes in the nature of the fixed effective connectivity involved the CRBL. Indeed, the closed loop between CRBL and SMAPMC was excitatory in AE and inhibitory in AO. Moreover, the extrinsic connectivity from the CRBL to the driving region V1 was inhibitory in AE and excitatory in AO.

### 2.2 Modulation of fixed effective connectivity (Step2)

Different modulations on the fixed effective connectivity between regions of the visuo-to-plan loop were demonstrated in AE and AO. The CRBL-SMAPMC loop was central in influencing motor planning with non-linear effects in AE but not in AO (Figure 2). For the analysis, this loop was split into two components: a forward pathway from V1 to CC and a backward pathway from CC to V1 (Figure 2).

**Figure 2.**
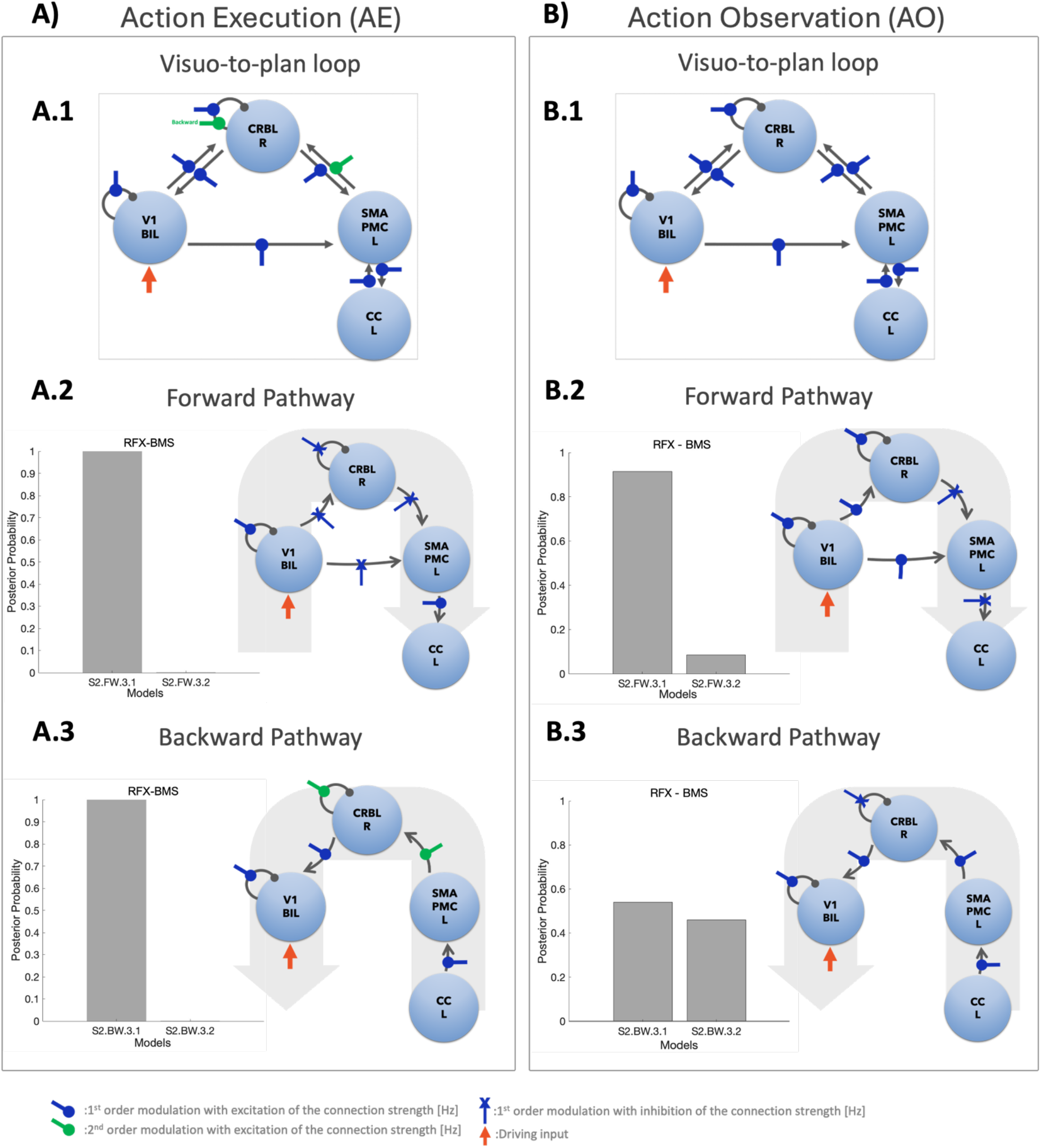
Modulation of the fixed effective connectivity. Forward and backward winning models are shown both for the Action Execution (AE) and the Action Observation (AO) conditions (Panel **A** and **B**, respectively). Each frame shows the posterior probability resulting from Random Effects Bayesian Model selection (RFX-BMS) (histogram) and a schematic representation of the S2-winning model. **A1)** AE S2-winning model showing the modulations on visuo-to-plan loop. **A2)** Forward Loop: RFX-BMS identifies the S2-forward winning model with a probability >90%. The signal linearly propagates from V1 to CC via SMAPMC, with an increased V1 self-connection and linear modulation of the SMAPMC-CC connection. Modulations on V1-CRBL, V1-SPMAPMC, CRBL-SPMAPMC and CRBL self-connectivity decrease the effective connectivity strength (blue cross-head arrow = inhibitory linear modulation). **A3)** Backward Loop: RFX-BMS identifies the S2-backward winning model with a probability >90%. Modulations are excitatory, increasing the strength of the effective connectivity. Modulations on SMAPMC-CRBL and CRBL self-connection resulted non-linear (green round-head arrow = excitatory non-linear modulation). **B1)** AO S2-winning model showing the modulations on visuo-to-plan loop. **B2)** Forward Loop: RFX-BMS identifies the S2-forward winning model with a probability >90%. Linear excitatory modulations characterise V1-CRBL and V1-SMAPMC connections (blue round-head arrow = excitatory linear modulation). Self-connections (V1 and CRBL) are increased by linear modulation. Inhibitory effects characterise the connections CRBL-SMAPMC and SMAPMC-CC. **B3)** Backward Loop: RFX-BMS identifies the S2-backward winning model with a probability of almost 60%. From CC to V1 the modulation is positive and linear. CRBL self-connection is linear and inhibited. V1 = bilateral primary visual cortex, M1 = left primary motor cortex, SMAPMC = left supplementary motor and premotor cortex, CC = left cingulate cortex, SPL = left superior parietal lobule, CRBL = right cerebellum.

#### Action Execution

RFX-BMS selected the winning model of each family with a probability higher than 70%, with an overall winning model identified with a probability of >90% (Figure 2A).

The winning model provided the best explanation of how BOLD-GF relations were modulated from one region to another in the visuo-to-plan loop (Figure 2A.1). The Step2 winning model showed an inhibitory linear modulation in the forward pathway from V1 to CRBL (Figure 2A.2), from V1 to SMAPMC as well as from CRBL to SMAPMC.

SMAPMC instead had a linear excitatory effect on the fixed effective connectivity strength between CC and SMAPMC, bidirectionally.

Non-linear modulatory effects were detected along the backward pathway, including the following connections: from SMAPMC to CRBL as well as within the CRBL self-connection (Figure 2A.3).

#### Action Observation

RFX-BMS selected the winning model of each family with a probability higher than 60%, with an overall winning model identified with a probability of >90% for the forward and 60% for the backward pathway (Figure 2B).

The winning model showed that the BOLD-GF relation was modulated linearly from one region to another (Figure 2B.1). In the visuo-to-plan forward pathway, CRBL turned excitation coming from V1 into inhibition towards SMAPMC and CC (Figure 2B.2). The backward pathway resulted in a fully excitatory modulation from one region to another with the only exception in the CRBL self-connectivity (Figure 2B.3).

Details of each family of models included in these analysis as well as results of RFX-BMS are reported in the supplementary material (Figures Sup.1-Sup.4).

## 3 Discussion

This work demonstrates the causal interaction between the CRBL and the cerebral cortex in two different functional conditions entailing the execution or observation of an action (AE or AO). The CRBL turns out to play a key role in shaping the effective connectivity of the loop ^12^ and to control it differentially depending on whether functional activation happens because of executing or observing a task.

Previous work looked at regional responses to task performance determining functional connectivity without questioning causality^5,14^. It is well known that effective connectivity, as quantified by DCM, yields asymmetric connections that allows to explore not only the strength of the effective connectivity but also its inhibitory and excitatory nature. Indeed, the effective connectivity of the motor network unveiled patterns characterized by an excitatory response in AE and an inhibitory response in AO. Moreover, modulations demonstrated a linear transformation of the BOLD-GF relation across the visuo-to-plan forward pathways, and non-linear transformations in the backward pathway between SMAPMC and the CRBL and within the CRBL self-connectivity only in AE. The interaction between these regions showed that there is a further elaboration of the BOLD-GF response, which changes the complexity of the polynomial signal function (e.g., from 1^st^ order to 2^nd^ order polynomial).

Investigating linearities and non-linearities is particularly interesting, since it can disentangle the role of each region activated during a task. We can interpret this as the CRBL working in predictive mode in AE, where the forward connection between the CRBL and SMAPMC remains linear to transfer an updated motor plan to cortical regions. This is different in AO, where the CRBL self-excitation or self-inhibition are reversed and all modulations are linear, possibly because of a lack of sensory feedback, as discussed in detail below. Overall, the emerging role of the CRBL is consistent with the theory that the CRBL uses internal models to adapt both motor actions and mental activities according to the context, operating as a forward controller that executes basic computational functions, namely timing, prediction, and sequence learning^1,15,16^.

### 3.1 Fixed Effective Connectivity (Step1)

Distinct regulatory mechanisms governing communication flow during AE and AO were revealed through the identification of contrasting excitatory and inhibitory connectivity patterns. Specifically, the cerebro-cerebellar motor planning circuit, encompassing the CRBL, SMAPMC, and the CC, showed increased activity in AE and a decreased activity in AO. This dichotomy occurs despite the loop being functionally engaged in both scenarios, highlighting the unique neuronal processes underlying each function. For instance, the cerebro-cerebellar motor planning loop (i.e., CRBL - SMAPMC - CC) exhibited excitation in AE but inhibition in AO; this is in line with the concept that the motor plan needed to perform an action is facilitated by the CRBL excitation of SMAPMC^1,17^.On the other hand, during observation there is no need for a motor plan, hence the CRBL inhibits SMAPMC. From a microscale perspective, this indicates that there is a different recruitment of neuronal populations depending on executing or only observing an action. While this information is available through DCM analysis, microscale heterogeneity linking functional activity to excitatory or inhibitory neurons may not be taken into consideration in macroscale connectivity studies; our results, indeed, supports the need for multi-scale approaches in line with previous literature^8,18,19^.

### 3.2 Modulation on Fixed Effective Connectivity (Step2)

The communication flow was extensively investigated to understand how the BOLD-GF modulation propagated between regions. Our key result is that non-linear modulations on the fixed effective connectivity were detected only in AE. This suggests that in the AE forward pathway, SMAPMC receives signals linearly from V1 and the motor plan is therefore initiated; SMAPMC then sends back signals non-linearly to the CRBL that also receives signals from V1, linearly. The emergent finding of a cerebellar non-linear self-connectivity may reflect the role of the CRBL as a forward controller in AE. Indeed, as summarized in figure 2A.1, a plausible scenario is that the CRBL is activated by contextual information coming from the visual system that provides feedback about AE; it activates internal inverse models that are compared to the motor plan received from SMAPMC; it then corrects the plan and feeds it back to SMAPMC for a continuous update of the motor plan ^8^. In other words, the CRBL receives a continuous input from V1 about the AE feedback. V1 also sends signals to SMAPMC that makes the best guess about the motor plan and sends it to the CRBL, non-linearly. The CRBL compares the two, performs error detection and internally updates the plan with motor prediction. It then sends the updated plan to SMAPMC, linearly.

At the microscale level, these non-linear modulations on the fixed effective connectivity are probably originating from the neuronal activity of SMAPMC, with bursting for motor control and/or adaptation for motor learning. Moreover, the non-linear modulation on CRBL self-connectivity could be the result of a continuous update of the motor plan by the CRBL itself. This interpretation comes from computational models of the CRBL where it has been shown that, e.g., the burst and pause patterns of Purkinje Cells are non-linear with respect to their input and are essential for learning and for error processing ^20,21^. From experimental models, we also know that there are non-linear relations between the frequency of excitation of cerebellar mossy fibers and the vasodilation that is at the basis of local neurovascular coupling ^7^

During AO, instead, modulation of fixed effective connectivity between or within regions was always linear, reflecting that predictive/control functions are less required^1,20^. As motor planning and action execution are not needed in AO, it can be suggested that the SMAPMC and the CRBL, are not engaged in sensorimotor transformations and linearly transfer signals coming from V1 throughout the network. This difference in how the CRBL modulates fixed effective connectivity between AE and AO is consistent with Schmahmann et al.^22^, who suggested that the general role of the cerebellum is to modulate behaviours around a homeostatic baseline to make them accurate and context-appropriate.

### 3.3 Future perspectives

Understanding the non-linear dynamics of the neuronal activity in SMAPMC and the CRBL is crucial for gaining insights into the CRBL as a forward controller in large-scale networks, driving motor planning and adaptation^23,24^. Our work points out the need to always include cerebellar nodes when studying AE and AO tasks, to avoid the risk of missing details that are critical to understand brain dynamics. This is particularly relevant in studies of neurological conditions characterized by motor impairment, from cerebellar ataxia to neurodegenerative or neuroinflammatory diseases. As an example, it has been shown that alterations in synaptic connectivity and neural adaptation could contribute to differences in the functional response to gripping tasks performed by MS patients in comparison to healthy controls^25– 27^. Moreover, it was shown that the BOLD-GF relation, using the same AE visuomotor task of this study, was altered in the posterior Brodmann area 4 of MS patients compared to healthy controls^6^. Given that MS patients fail in sensorimotor integration due to demyelination and neurodegenerative processes, linear and non-linear modulations on fixed effective connectivity recorded during AE may be affected. An interesting hypothesis could be to study the cerebellar non-linearities in MS as the CRBL may fail to function in predictive and execution modes. Understanding the linear/non-linear modulations of BOLD signals in health and disease could be pivotal for efficient design of forward controllers that may be used for rehabilitation strategies designed to enhance neural adaptation and improve motor function ^28,29^.

### 3.4 Study Considerations

DCM has been valuable for advancing the understanding of the causality at the origin of brain interactions^30–32^. Nevertheless, DCM is a data-driven formalism as its Bayesian model inversion depends upon the data at hand^33^. In the AO paradigm, a smaller cohort was included (see Methods section), which resulted in a discrepancy in sample size compared to the AE paradigm. To address any potential bias arising from this difference in cohort sizes, Bayesian model selection was performed using random-effects analysis^13^. Furthermore, models tested required a compromise between the number of regions and connections included and the computational resources. Multiple model configurations focusing on pathways (i.e., forward and backward pathways from V1) were tested to efficiently estimate microcircuitry parameters and to prevent model overfitting and computational demands.

DCM uses a Taylor approximation to neuronal dynamics as a generative model, which is based on theoretical assumptions rather than biological evidence such as the structural and functional differences of cortical and subcortical regions^11^. First attempts to introduce region specific models have been implemented with promising results for the cortex, such as the model of the canonical microcircuit, as well as for subcortical regions^21,36,37^. Amongst these, a recently developed cerebellar mean field model could be used within the DCM framework for extracting more physiologically grounded estimations of the neural basis of the BOLD non-linearity propagations ^21^.

A further consideration must concern the fact that the CRBL works as a forward controller not only in motor function but also in cognition. The task used for this study of causality between regions involves the motor network, but also a broader range of associative regions ^5,14^, not included in this DCM analysis for the reasons described above. It is possible to hypothesise that, given the modular cerebellar microstructure, the CRBL exerts a similar role in modulating fixed effective connectivity in networks supporting cognitive tasks, and that it is implicated in defining the nature and modulation of the fixed effective connectivity between regions, affecting a broader spectrum of functions.

### 3.5 Conclusions

DCM analysis demonstrates that executing and observing an action is supported by the same cerebro-cerebellar network, whose modulation discriminates AE from AO. The difference between these tasks is that the first engages sensorimotor interactions, while the second involves an internal simulation process. While the CRBL is well positioned in both cases to operate as the generalized forward controller as predicted by theory^2,3^, an important addition of this work is that the CRBL can dynamically change its functional mode depending on the task. The underlying circuit mechanisms remain to be determined, though. Physiologically, the excitatory/inhibitory switch from AE to AO could reflect a different balance between excitation and inhibition in the deep cerebellar nuclei (DCN), that are modulated by excitatory mossy fibre collaterals and inhibitory Purkinje cell axons^38^.The DCN then project to thalamic nuclei, which in turn project to the cerebral cortex^39^. Then, the cerebellar regulation of SMAPMC could reflect changes in gamma-band coherence between cortical areas, as recently revealed in mice^40^. Finally, signals conveyed to the CRBL through sensory pathways during AE may set up non-linear dynamics in the cerebellar internal microcircuit^9,41^ as well as in other parts of the extended visuomotor network. Future work is warranted to further understand the neural underpinning of the present observations, and to extend the present results in the context of neurological diseases and sensorimotor/cognitive robotic controllers.

## Methods

### Subjects

This analysis was performed on subjects already included in previous investigations^5,14^. Twenty-one healthy volunteers (aged 22 ± 4 years, 9 males) were enrolled for the action execution (AE) paradigm, while nine healthy volunteers (aged 26 ± 8 years, 3 males) were included in the action observation (AO) one. The participants did not have any history of neurological or psychiatric diseases. All the participants received a detailed explanation of the experimental procedures before participating in the experiment. The study was approved by ethics committee and all participants gave their written informed consent^5,14^.

### MRI acquisitions

A 3D-T1 weighted (3DT1) MPRAGE volume (TR/TE/TI = 6.9/3.1/824 ms, flip angle = 8°, voxel size = 1×1×1 mm) and three T2*-weighted GE-EPI fMRI series (TR/TE = 2500/35 ms, flip angle = 90°, voxel size = 3×3×3 mm, 200 volumes) were acquired on a 3T Philips Achieva scanner (Philips Healthcare, Best, The Netherlands) with a 32-channel head coil

### Experimental Design

The paradigm involved an event-related power grip task using an fMRI-compatible squeeze ball, as previously described in Alahmadi et al., 2016. Participants engaged in a visuomotor execution task (i.e., an AE task) and an observation task (i.e., an AO task). The AE task consisted of a total of 75 active trials, evenly distributed across 5 different Grip Forces (20%, 30%, 40%, 50%, 60% of each subject’s maximum voluntary contraction), performed by each participant. The AO task involved watching a video that displayed the right hand of an actor performing the squeeze ball task, without any indication of the applied force.

### fMRI preprocessing

fMRI images were pre-processed with a customized MATLAB v2019b script combining commands of different tools, namely SPM12 (https://www.fil.ion.ucl.ac.uk/spm/), FSL (https://fsl.fmrib.ox.ac.uk/fsl/fslwiki) and MRtrix3 (https://www.mrtrix.org/). The steps of the preprocessing included the intensity correction, the Marchenko-Pastur principal component analysis (MP - PCA) denoising^42^, slice timing correction, realignment to the mean functional volume; 3DT1 images were normalized to the standard Montreal Neurological Institute (MNI) space and affine registration of fMRI images were carried out. Then, polynomial detrending of the signal was applied to remove cerebro-spinal fluid(CSF) contributions as identified by a CSF region of interest placed in the ventricles and to correct for motion using 24 motion regressors obtained from the registration^43^.

### Time-series extraction

The pipeline used to extract time-series in specific volume of interest (VOIs) included in the motor network is shown in Figure 3 and the technical specifications are reported in Table 1 for AE and Table 2 for AO. For the sake of clarity, in figures we report the acronym of each region also indicating the laterality (Left (L), Right (R) and Bilateral (BIL); e.g. Right CRBL is reported as CRBL R). For each subject, a mask was defined for each of the brain regions embedded in the network by implementing a conjunction analysis with anatomical and functional constraints. The anatomical constraints were defined by an anatomical parcellation computed by merging a cerebral and a cerebellar atlases, respectively the Brodmann^44^ and a spatially unbiased atlas template of the cerebellum and brain stem (SUIT) ^45^ atlases (Figure 3A). The functional constraints, instead, were defined with a second-level analysis, which computed the group activation peak (GAP) in each activated region (Figure 3B). Then, a geometrical mask centred on subject-specific activation peaks and including only the activated voxels was restricted to the anatomical and functional constraints (Figure 3C). The dimension of the geometrical mask is defined accounting for the anatomical dimension of each VOI. The same procedure was used to extract each region. Significance thresholds were defined according to the region and the nature of task (AE in Table 1 and AO in Table 2). A time-series was finally extracted for each VOI as the average of the functional signals of the voxels belonging to the geometrical mask.

**Table 1.**
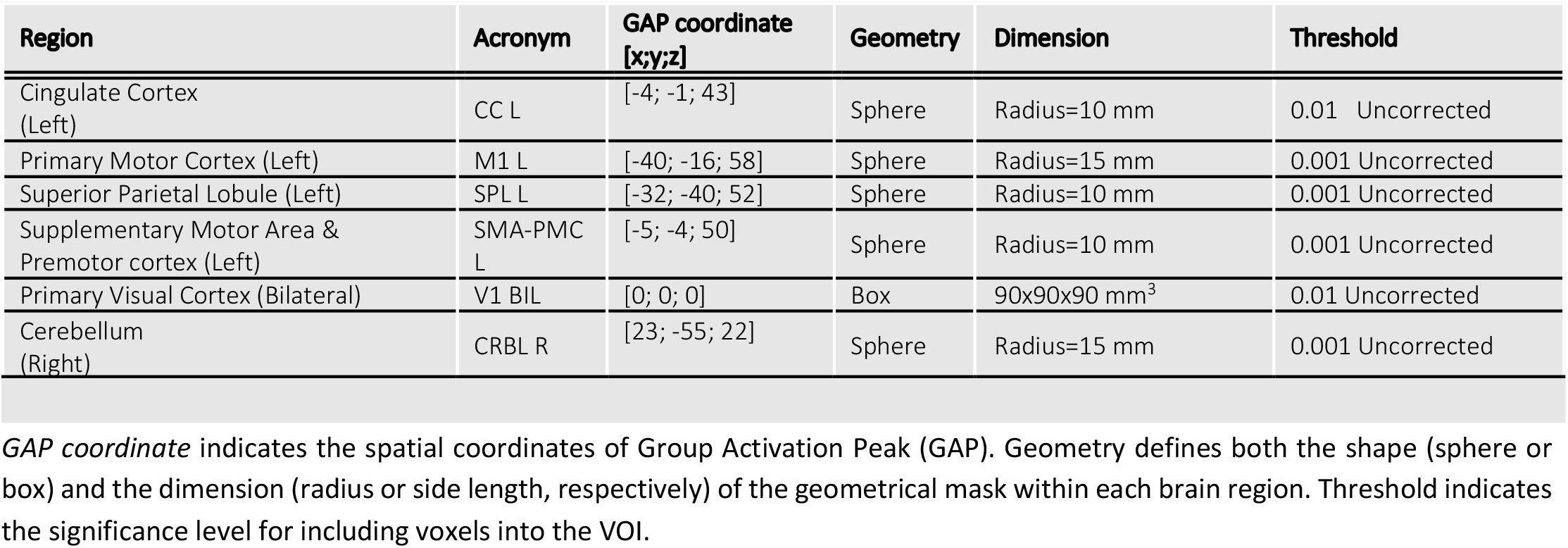
Details for Volume of Interest in the Action Execution task.

**Table 2.**
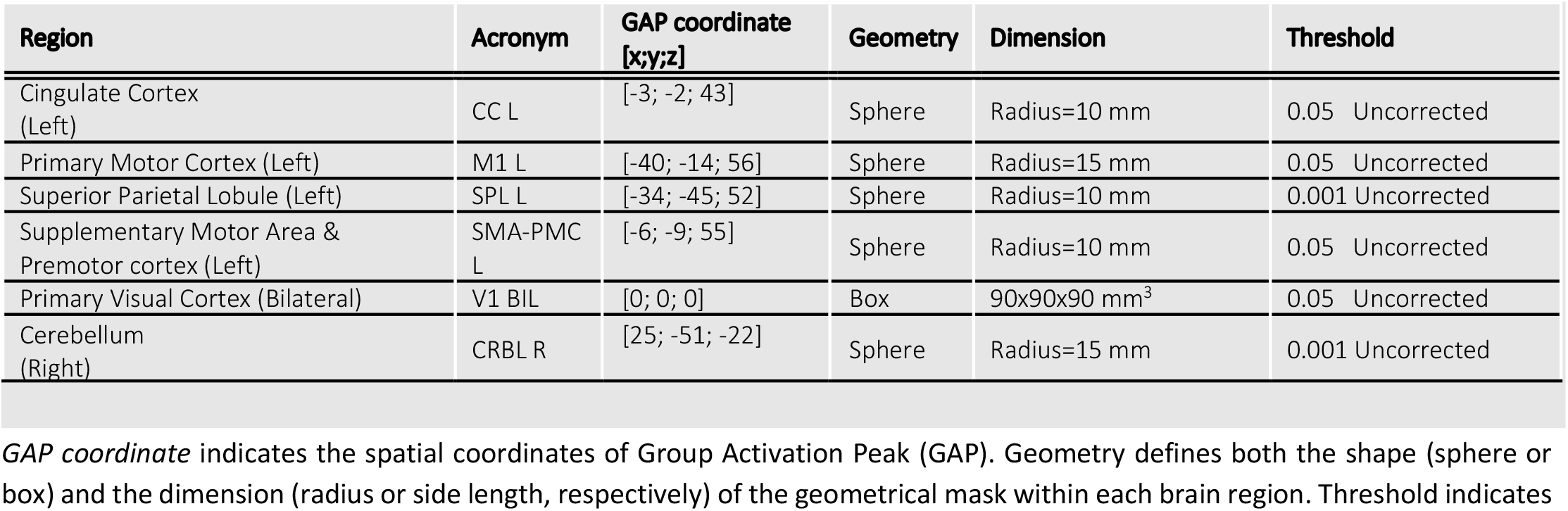
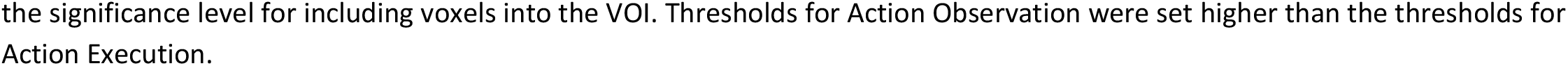
Details for Volume of Interest in the Action Observation task.

| Region | Acronym | GAP coordinate [x;y;z] | Geometry | Dimension | Threshold |
| --- | --- | --- | --- | --- | --- |
| Cingulate Cortex (Left) | CC L | [-3; -2; 43] | Sphere | Radius=10 mm | 0.05 Uncorrected |
| Primary Motor Cortex (Left) | M1 L | [-40; -14; 56] | Sphere | Radius=15 mm | 0.05 Uncorrected |
| Superior Parietal Lobule (Left) | SPL L | [-34; -45; 52] | Sphere | Radius=10 mm | 0.001 Uncorrected |
| Supplementary Motor Area & Premotor cortex (Left) | SMA-PMC L | [-6; -9; 55] | Sphere | Radius=10 mm | 0.05 Uncorrected |
| Primary Visual Cortex (Bilateral) | V1 BIL | [0; 0; 0] | Box | 90x90x90 mm<sup>3</sup> | 0.05 Uncorrected |
| Cerebellum (Right) | CRBL R | [25; -51; -22] | Sphere | Radius=15 mm | 0.001 Uncorrected |

*GAP coordinate* indicates the spatial coordinates of Group Activation Peak (GAP). Geometry defines both the shape (sphere or box) and the dimension (radius or side length, respectively) of the geometrical mask within each brain region. Threshold indicates
the significance level for including voxels into the VOI. Thresholds for Action Observation were set higher than the thresholds for Action Execution.

**Figure 3.**
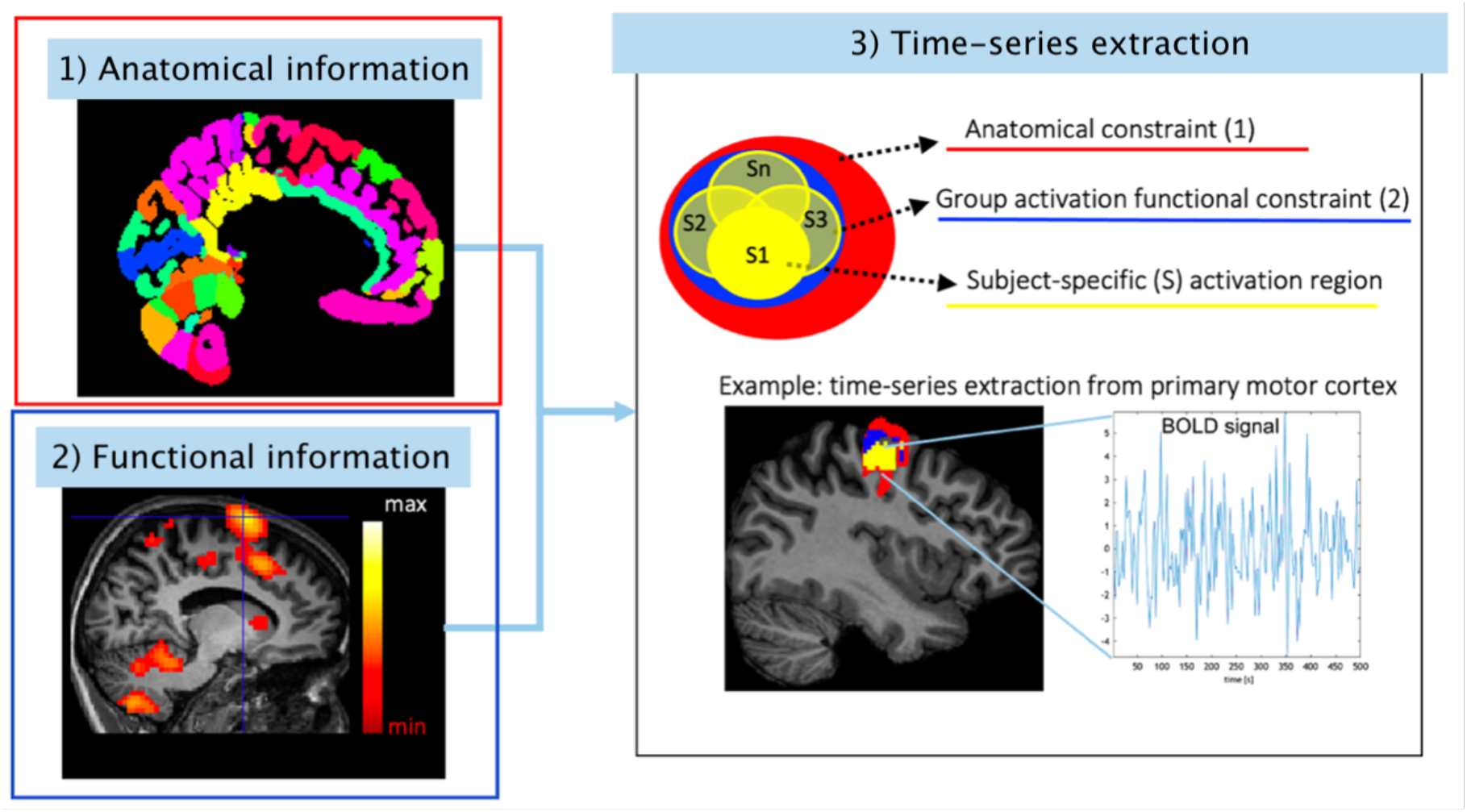
Pipeline to extract fMRI time-series. For each region of the motor network, the anatomical mask is extracted from an ad-hoc atlas combining the Brodmann and the spatially unbiased atlas template of the cerebellum and brainstem (SUIT) atlases **(1)** and the group activation peak is computed **(2)**. Subject-specific time-series were extracted in a VOI obtained restricting a geometrical mask to the anatomical and functional constraints **(3)**.

### Dynamic causal modelling

DCM analysis using DCM version 12.5 implemented in SPM12 was performed to infer the effective connectivity of the motor network both in AE and AO conditions.

Effective connectivity was computed accordingly to^46^:

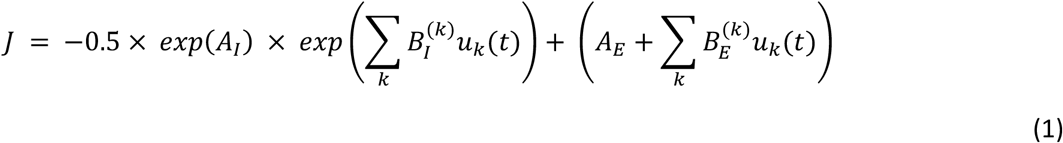

Where ***J*** is the estimated effective connectivity, ***A***_***I***_ is the self-connectivity (i.e., regions intrinsic connectivity) which is unitless representing a logarithmic scaling parameter and multiplied by the default value of -0.5 Hz, ***A***_***E***_ is the extrinsic effective connectivity (Hz), ***B***_***I***_ and ***B***_***E***_ are, respectively, the modulations on the intrinsic and extrinsic effective connectivity of the *k*^*th*^ region, which are driven by the external input ***u****(t)*.

Specifically, DCM analysis was implemented to provide firstly, the fixed effective connectivity of the motor network, quantified by estimating the ***A***_***I***_ and ***A***_***E***_ matrices (S1, Figure 4). Then, after the best model for effective connectivity was identified, GF modulatory effects in the forward and backward visuo-to-plan pathways was assessed (S2, Figure3) by estimating the values of B (i.e., **B**_***I***_ and **B**_***EI***_) matrices.

### Model constructions

We conducted the analysis in two levels:

- Step 1 (S1): to estimate the strength of the connections between the regions of the motor network, namely matrix **A** defined as *fixed* effective connectivity (Figure 4);
- Step 2 (S2): to quantify the strength of the task-dependent coupling between regions namely matrix **B** modelling the modulation on the fixed effective connectivity (Figure 5).

**Figure 4.**
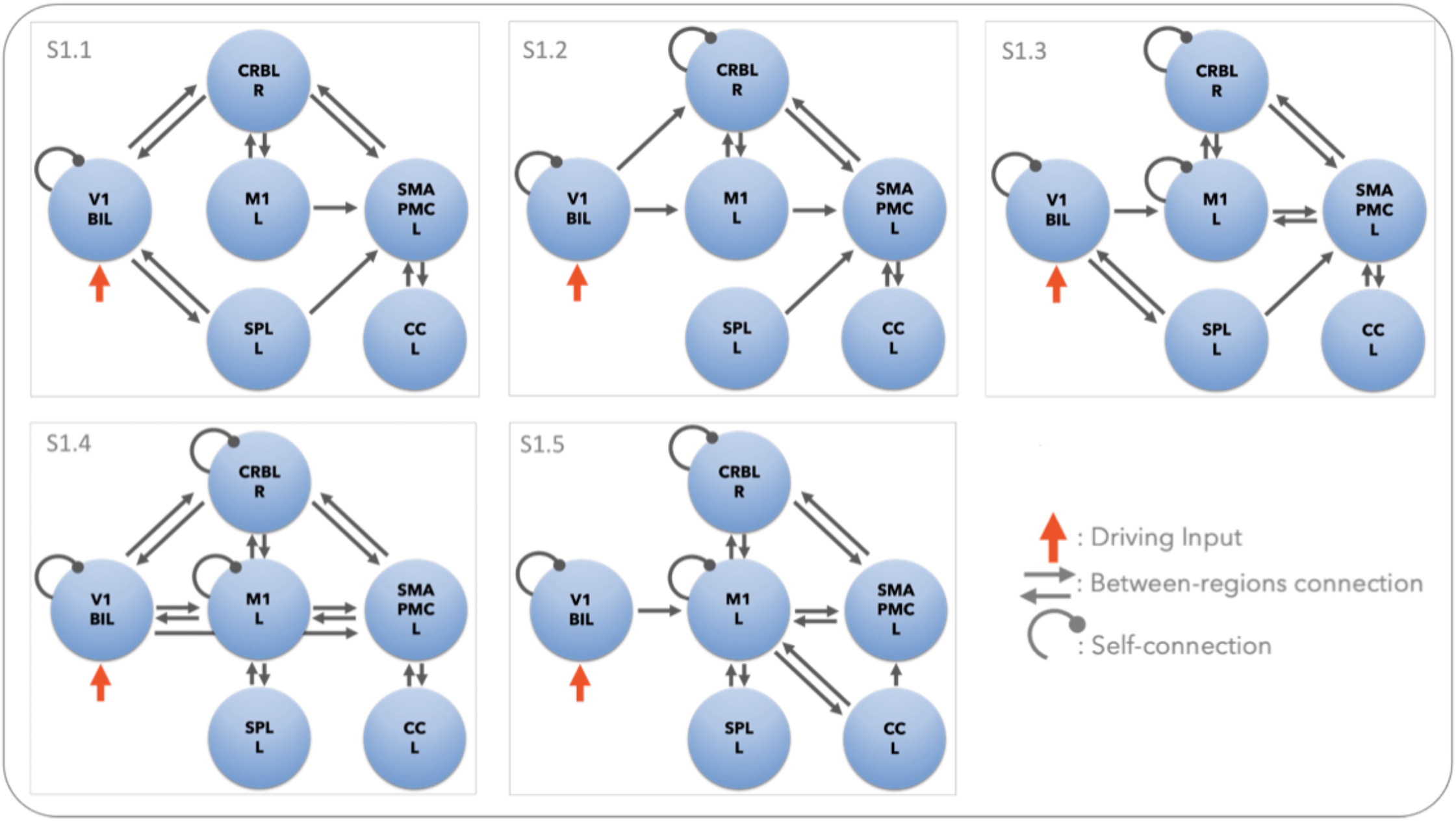
Models to assess the fixed effective connectivity. Schematic illustration of the fixed effective connectivity models (S1.1 – S1.5) of cortico-cerebellar loops (step 1). The same configurations were tested for Action Execution (AE) and Action Observation (AO) separately. All models are driven by V1 given the visuomotor nature of the task (purple arrow). The directionality of between-region connections directionality was modelled to explore different configurations of the network causal relations. Self-connections, instead, allow us to investigate the sensitivity of a region to the network input. The strength of each effective connection is quantified in Hz by the Bayesian model inversion. V1 = bilateral primary visual cortex, M1 = left primary motor cortex, SMAPMC = left supplementary motor and premotor cortex, CC = left cingulate cortex, SPL = left superior parietal lobule, CRBL = right cerebellum.

**Figure 5.**
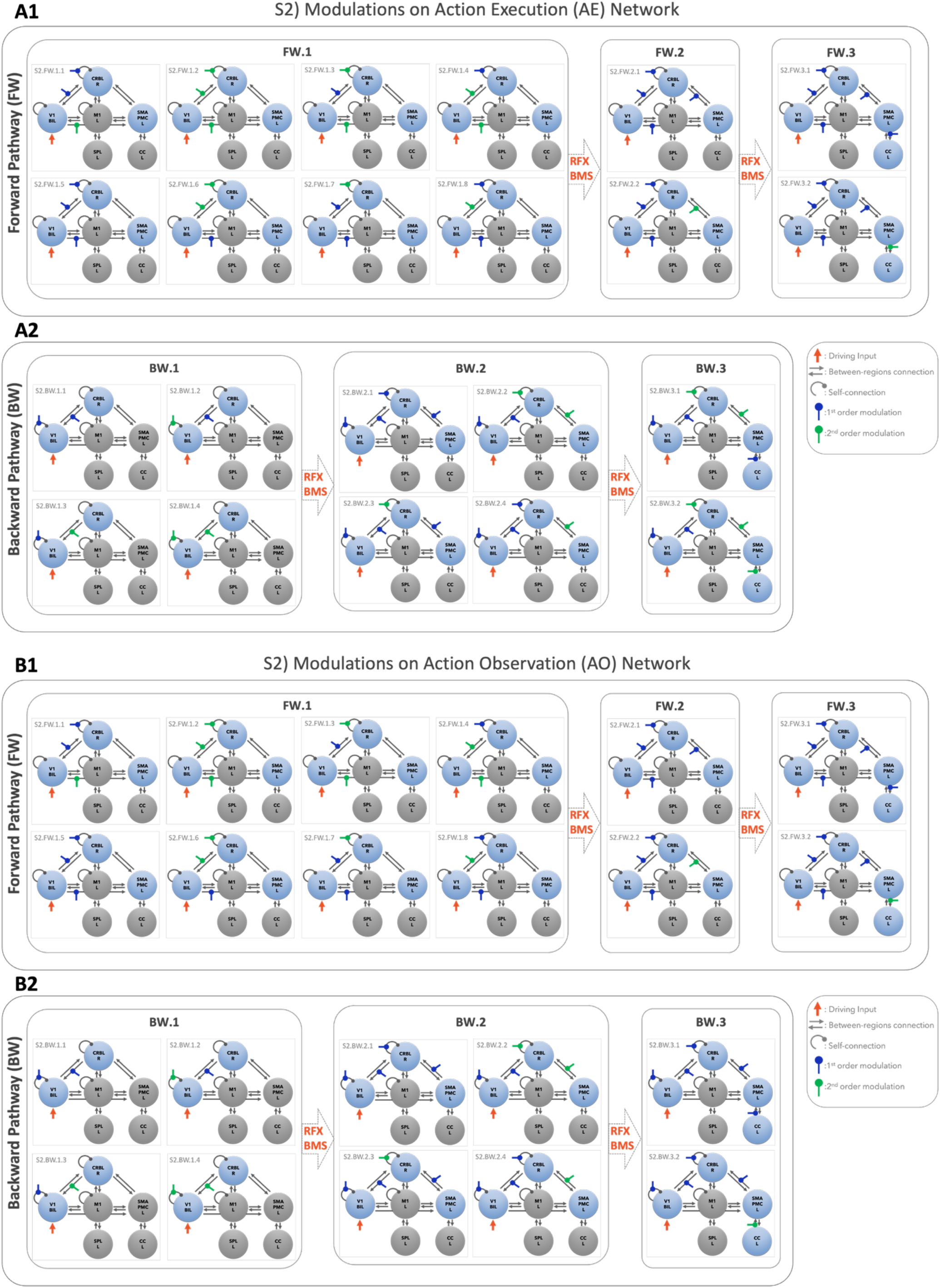
Models to assess possible modulations on the effective connectivity. Schematic illustration of modulation testing on Action Execution (AE – Panel A1, A2) and Action observation (AO – Panel B1, B2) effective connectivity models of cortico-cerebellar loops (Step 2). As effective connectivity is directional, it was possible to define forward and backward pathways. Forward pathways (FW) were defined from V1 to CRBL, SMAPMC, and CC (Panel A1 and B1). Backward pathways (BW) were defined from CRBL, SMAPMC and CC to V1 (Panel A2 and B2). For each effective directional connection between pairs of regions, we defined possible modulatory effects of the BOLD signal from the starting region to the target region. Different combinations of linear/non-linear modulations were tested. Random Effects Bayesian Model selection (RFX-BMS) was performed to identify the winning model at each stage of the stack procedure. A stack procedure grouping different modulations of the same connection was applied: the output of the simpler configuration (Family i) become the input for the more complex configurations (Family i+1), with i= index of Family. For both backward and forward pathways, the winning model of the most complex was the overall winning model (i.e., Family FW.3 and BW.3 for forward and backward pathway respectively). V1 = bilateral primary visual cortex, M1 = left primary motor cortex, SMAPMC = left supplementary motor and premotor cortex, CC = left cingulate cortex, SPL = left superior parietal lobule, CRBL = right cerebellum.

#### S1) Fixed effective connectivity

By the nature of the task, V1 was defined as the driving region (*u(t)*) triggering the activity of the network in all the hypotheses. Based on prior knowledge, a set of five models with different configurations of inward and outward connections were defined to quantify the effective connectivity of the AE and AO conditions (Figure 4)^47–49^.

Self-connections were explicitly included for the driving region V1 in all S1 models, and for CRBL for S1 models that investigate the changes in cerebellar neuronal activity triggered by the forward and backward loop with M1 and SMAPMC (Figure 4 – models S1.2 to S1.5). Focusing on the visuo-to-plan pathway, bidirectional connections between SMAPMC and CRBL were included to all S1 models.

In practice, model construction was defined following the standard procedure described by Zeidmann et al., 2019^46^: ***A***_***E***_*(k)* and ***A***_***I***_*(k)* were set to 1 Hz and 1, respectively, when the connection with a *k*^*th*^ region was included into the model, otherwise ***A***_***E***_*(k)* was set to 0 Hz when the specific connection was of no interest, thus preventing that the model inversion updated it. The same strategy was applied to ***A***_***I***_*(k)*, resulting in a prior value of –0.5 Hz (equation 1) meaning that it was updated by model inversion even if not explicitly included into the model. The difference between prior values of extrinsic and self-connections is due to the nature of what they model: indeed, extrinsic-connections model the inter-regions causal influence, while self-connections model the intra-region neuronal activity, which is always influenced by the network dynamics.

Specifically, different fixed effective connectivity models (S1.1 – S1.5) are based on literature review of structural connectivity and functional activation for similar tasks (Figure 4).

#### S2) Modulations on the effective connectivity

The effect of task-driven modulations of region-specific BOLD signals were tested starting from the effective connectivity model architecture, resulting from S1 (Figure 5). This analysis was focused on visuo-to-plan functional pathway embedding the loop between the visual input, cortical motor planning areas and the CRBL, considering single or bi-directional connections between pairs of regions, depending on the fixed effective connectivity resulting from S1-winning model. In details, first and second order polynomial expansions of the BOLD-GF relation as described in Alahmadi et al., 2016, were tested as modulatory effects on the visuo-to-plan functional loop including V1 - CRBL - SMAPMC – CC regions^5^. As effective connectivity is directional, it was possible to define forward and backward pathways, from V1 towards motor planning areas and from motor planning areas to V1 respectively. Self-connectivity is considered both in forward and backward pathways as inward connections (e.g., Figure 5A.1 in Model S2.FW.1.1 the forward pathway is defined from V1 to CRBL including the CRBL self-connection)

Various configurations are defined depending on the complexity of the regions included (see supplementary material for the RFX-BMS details). In Figure 5, forward and backward pathways are shown for three families of possible network configurations each, with increasing complexity from family 1 to family 3 (i.e. FW.1, FW.2, FW.3 and BW.1, BW.2, BW.3). For example, the simplest forward pathway starts from V1 and includes connections to CRBL and SMAPMC (Figure 5A1, Model S2.FW.1.1); the most complex forward loop starts from V1 and ends in CC, passing through CRBL and SMAPMC (Figure 5A1, Model S2.FW.3.2). The same rationale was used to define backward pathways (Figure 5B).

Following the same strategy as for the fixed effective connectivity model selection (S1), values in the B matrix were set to 1 Hz when the modulations were included in the model and to 0 Hz otherwise. A *stack* procedure of model evaluation was applied for both forward and backward pathways to AE and AO separately (Figure 5A and 5B respectively), increasing the complexity of the architecture, i.e., the number of modulations included, at each step The backward path was analysed following the same rationale.

### Model evaluation and statistics

Bayesian model inversion was implemented on AE and AO data to estimate the best models to explain BOLD signal responses from regions of the motor network, based on their fixed effective connectivity, namely matrix A, at a subject level^50^. Bayesian model selection (BMS) was used to select the best model for a different set of hypotheses at a group level^13^. For each model configuration, model inversion estimated the posterior probability of each connection (S1) and each modulation (S2) defined in the model construction, quantifying the model likelihood (i.e., model evidence) with the experimental data (i.e., fMRI time-series). Random Effects-BMS (RFX-BMS) was used to assess whether the best model (winning model) differed between subjects^13^. Bayesian Model Averaging (BMA) was used to provide coherent accounting of statistical structure by including weighted average of each connectivity strength over model space^50^. The S1-winning model was used as fixed architecture to set different configurations of the modulations (2.6.1 Model construction). RFX-BMS was applied at each step of the forward/backward path to find the S2-winnig model to define the propagation of the BOLD-GF modulations.

### Reproducibility of the framework

All the procedures for fMRI preprocessing, regions of interest selection and DCM analysis were run on a Desktop PC provided with AMD Ryzen 7 2700X CPU @ 2.16GHz with 32 GB RAM in Ubuntu 16.04.7 LTS (OS). The functions packages implemented to perform the selection of regions of interest and DCM analysis are available on https://github.com/RobertaMLo/CRBL_DCM.

