## Supplementary Material for "Cerebellar control over inter-regional excitatory/inhibitory dynamics discriminates execution from observation of an action"

### Action Execution (AE)

#### Forward Pathway

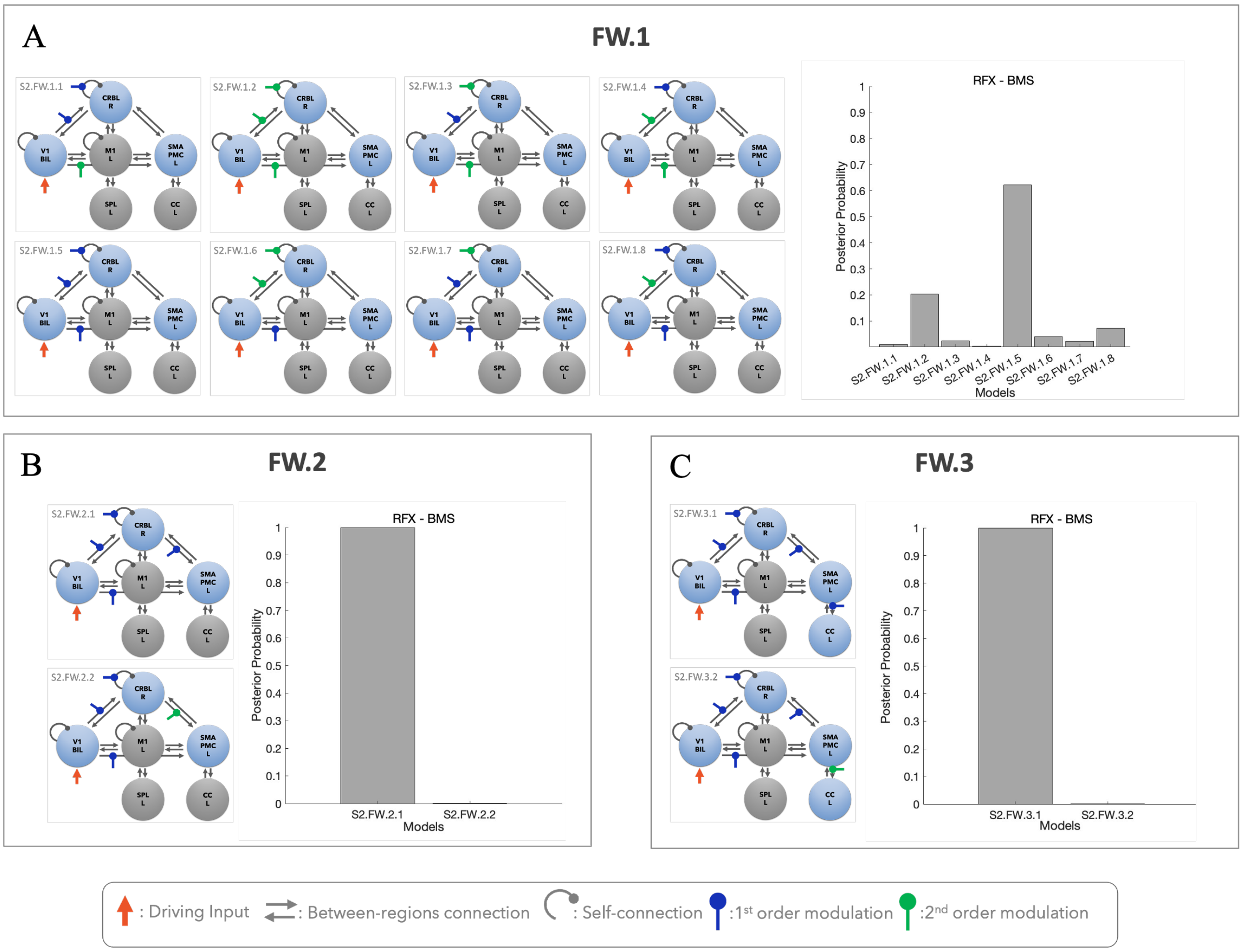


**Supplementary Figure 1 – AE Forward Pathway**

**A) Model family 1** of AE was constructed to investigate different configurations of linear/nonlinear modulations in motor planning anticipation triggered by the visual input including the connections from V1 to CRBL, CRBL self-connectivity and from V1 to SMAPC. RFX-BMS was applied to select the winning model.

**B) Model family 3** of AE investigate the cerebro-cerebellar interactions in motor planning on the winning model of the model family 1 by following a stack procedure: grouping different modulations of the same connections as applied in the winning model of the model family 1. Different configurations of CRBL-SMAPMC were applied and RFX-BMS was applied to select the winning model.

**C) Model family 5** of AE is the most complex model in the forward pathway, following the same stack procedure and including the modulation configurations to the winning model of model family 3, to investigate the associative area's interactions in motor planning including different modulation configurations from SMAPMC to CC. RFX-BMS was applied to select winning model. Winning model of the model family 5 is the overall winning model of the forward pathway network of AE.

### Action Execution (AE)

#### Backward Pathway

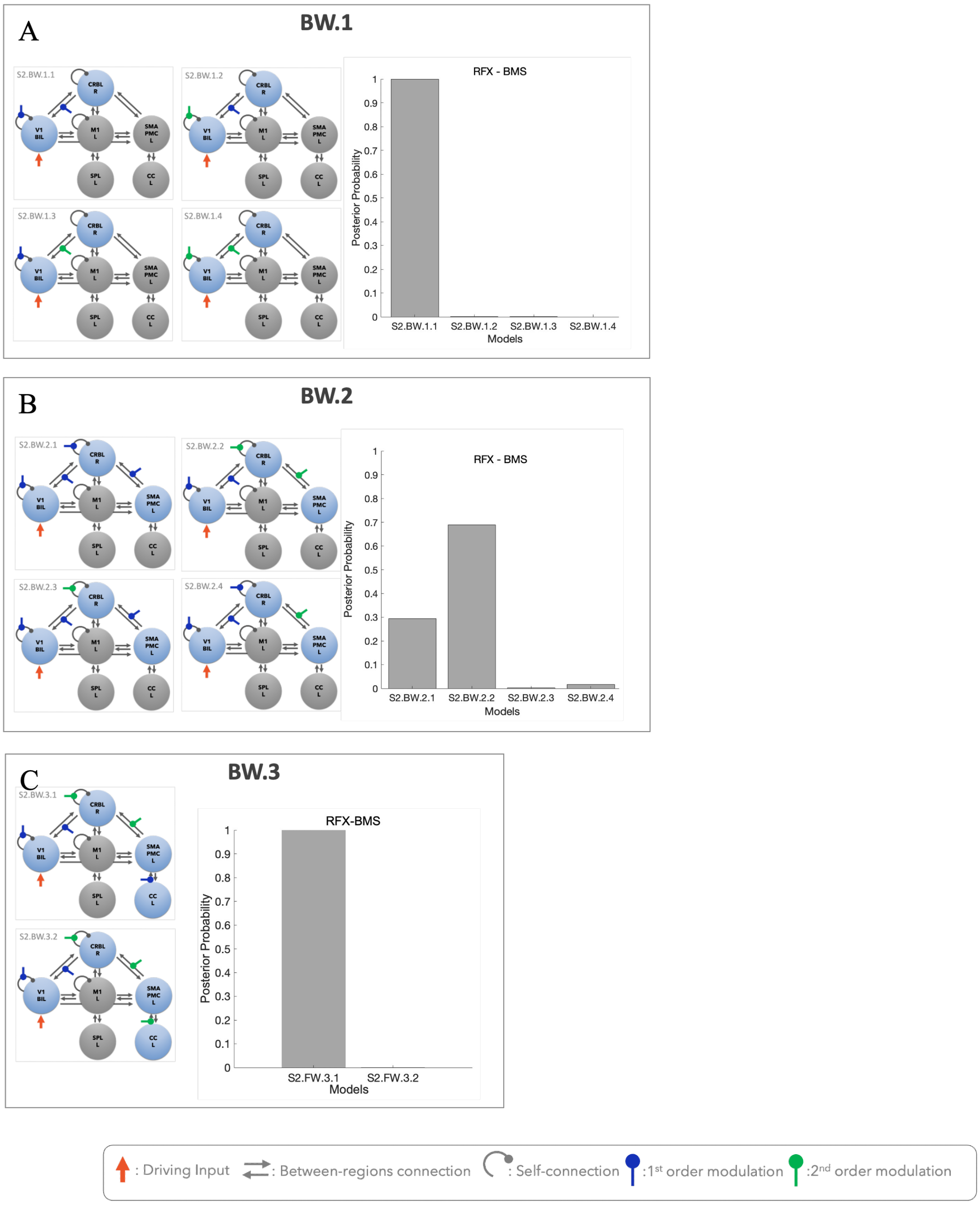


**Supplementary Figure 2 – AE Backward Pathway**

**A) Model family 2** of AE backward pathway was constructed to investigate different configurations of linear/nonlinear modulations in the feedback modulation in motor planning anticipation towards the visual input including the connections from CRBL to V1 and V1 self-connectivity. RFX-BMS was applied to select the winning model.

**B) Model family 4** of AE backward pathway, investigate the cerebro-cerebellar feedback in motor on the winning model of the model family 2 by following a stack procedure: grouping different modulations of the same connections as applied in the winning model of the model family 2. Different configurations of CRBL-SMAPMC and self-connectivity of the CRBL were constructed and RFX-BMS was applied to select the winning model.

**C) Model family 6** of AE backward pathway is constructed as the most complex model in the feedback pathway, following the same stack procedure and including the modulation configurations to the winning model of model family 4, to investigate the associative area's feedback in motor planning including different modulation configurations from CC to SMAPMC. RFX-BMS was applied to select winning model. Winning model of the model family 6 is the overall winning model of the backward pathway network of AE.

### Action Observation (AO)

#### Forward Pathway

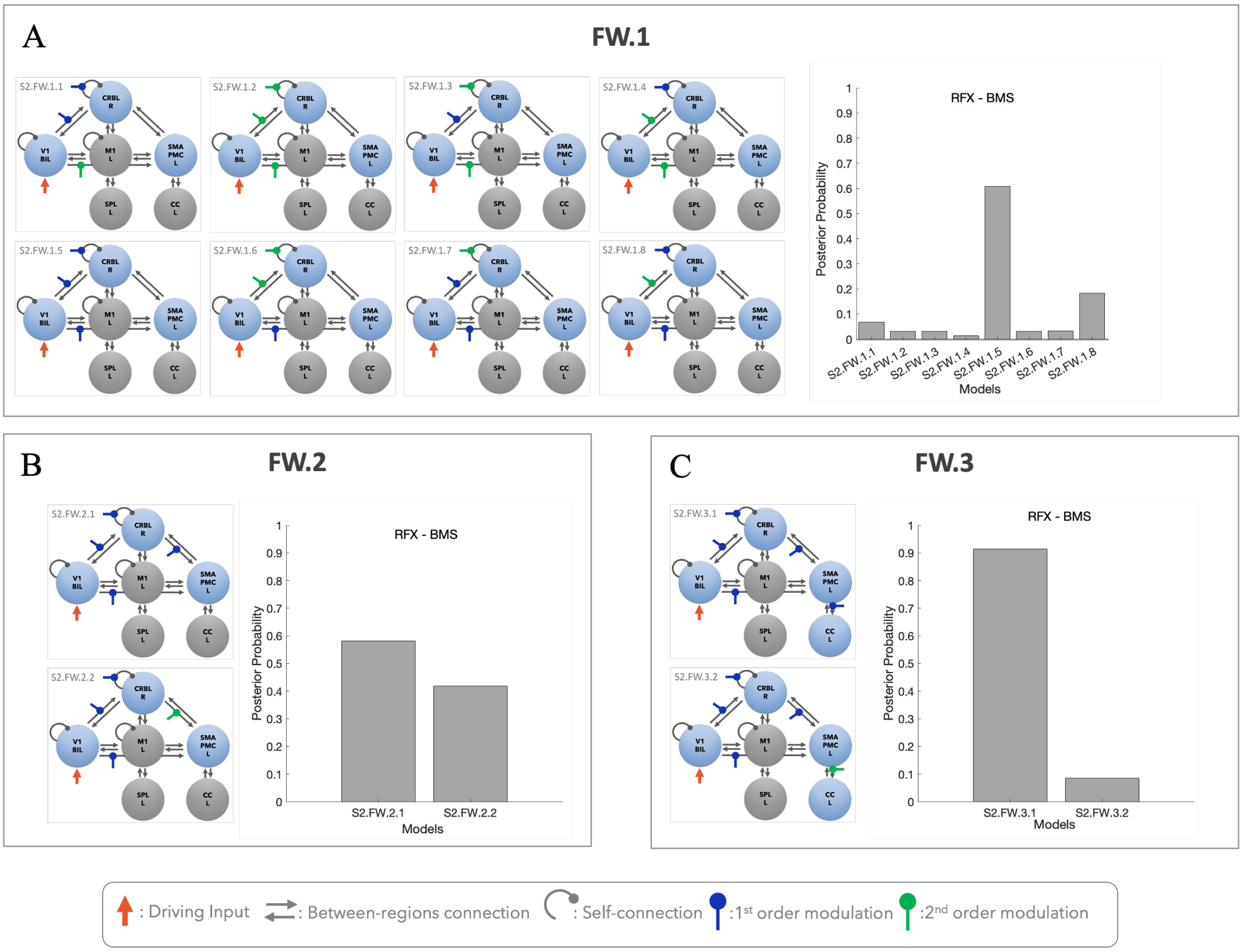


**Supplementary Figure 3 – AO Forward Pathway**

**A) Model family 1** of AO forward pathway was constructed to investigate different configurations of linear/nonlinear modulations in motor planning anticipation triggered by the visual input including the connections from V1 to CRBL, CRBL self- connectivity and from V1 to SMAPC. RFX-BMS was applied to select the winning model. Same model pipeline followed in AE forward pathway

**B) Model family 3** of AO is defined following the same pipeline for AE (see Supplementary Figure 1) cerebro-cerebellar interactions in motor planning are investigated on the winning model of the model family 1 by following a stack procedure: grouping different modulations of the same connections as applied in the winning model of the model family 1. Different configurations of CRBL-SMAPMC were applied and RFX-BMS was applied to select the winning model.

**C) Model family 5** of AO is the most complex model in the forward pathway, following the same stack procedure applied in AE and including the modulation configurations to the winning model of model family 3, to investigate the associative area's interactions in motor planning including different modulation configurations from SMAPMC to CC. RFX-BMS was applied to select winning model. Winning model of the model family 5 is the overall winning model of the forward pathway network of AO.

### Action Observation (A0)

#### Backward Pathway

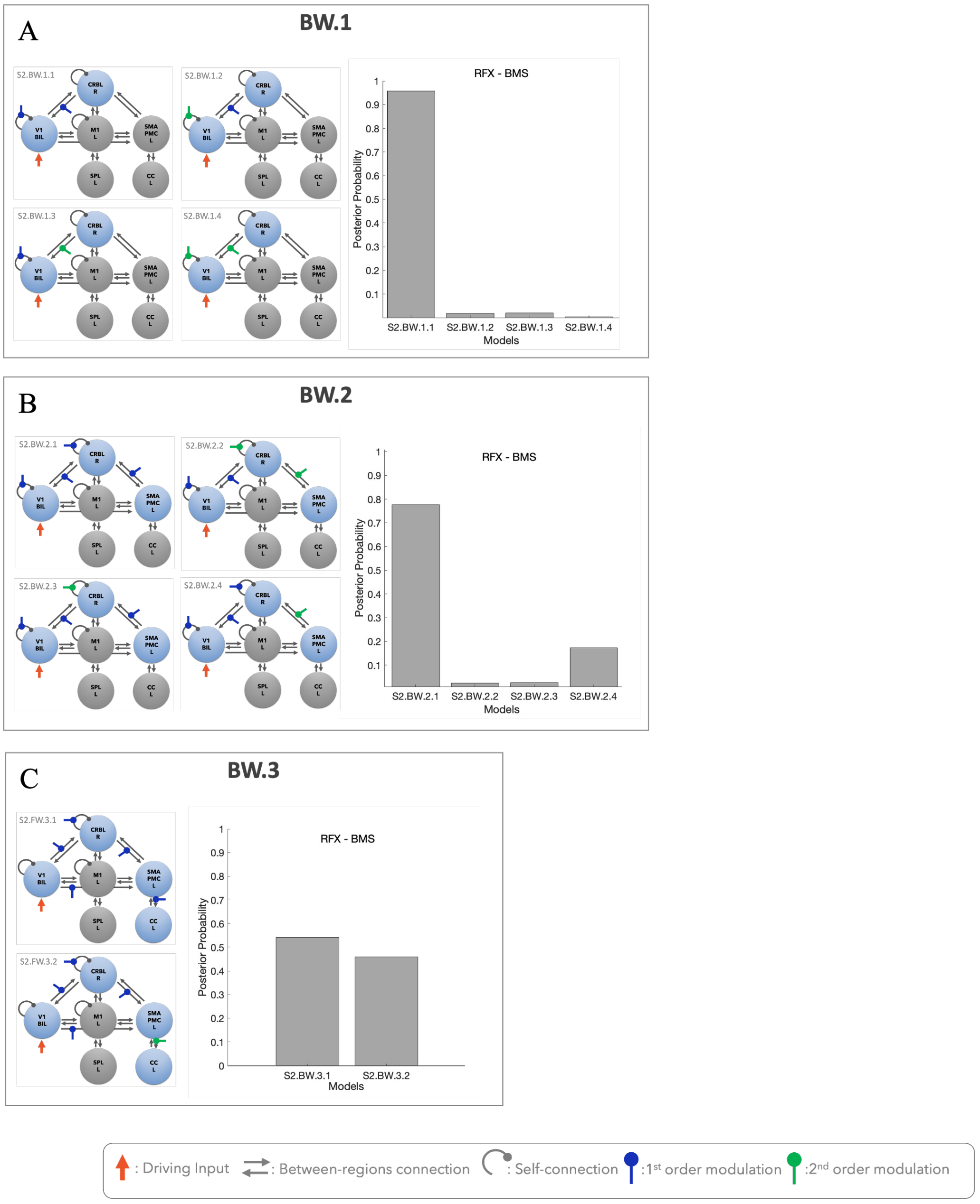


**Supplementary Figure 4 – AO Forward Pathway**

**A) Model family 2** of AO backward pathway was constructed to investigate different configurations of linear/nonlinear modulations in the feedback modulation in motor planning anticipation towards the visual input including the connections from CRBL to V1 and V1 self-connectivity. The same pipeline followed for AE was followed and RFX-BMS was applied to select the winning model.

**B) Model family 4** of AO backward pathway investigate the cerebro-cerebellar feedback of AO in motor planning on the winning model of the model family 2 by following a stack procedure followed in AE: grouping different modulations of the same connections as applied in the winning model of the model family 2. Different configurations of CRBL-SMAPMC and self-connectivity of the CRBL were constructed and RFX-BMS was applied to select the winning model.

**C) Model family 6** of AO backward pathway is constructed as the most complex model in the AO feedback pathway, following the same stack procedure of AE backward pathway and including the modulation configurations to the winning model of model family 4, to investigate the associative area's feedback of AO in motor planning including different modulation configurations from CC to SMAPMC. RFX-BMS was applied to select winning model. Winning model of the model family 6 is the overall winning model of the backward pathway network of AO.
